# The abundant-centre is not all that abundant: a comment to Osorio-Olvera et al. 2020

**DOI:** 10.1101/2020.02.27.968586

**Authors:** Tad Dallas, Samuel Pironon, Luca Santini

## Abstract

Species abundance is expected to decrease from the centre towards the edge of their ecological niches (abundant niche-centre hypothesis). Recently, Osorio-Olvera *et al*. (2020) reported strong support for the abundant niche-centre relationship in North American birds. We demonstrate here that methodological decisions strongly affected perceived support. Avoiding these issues casts doubt on conclusions by Osorio-Olvera et al. and the putative support for the abundant nichecentre hypothesis in North American birds.

---

The spatial distribution of abundance has long fascinated ecologists who searched for general rules governing where species occur and the density at which they are found (McGill *et al*., 2007; Sagarin & Gaines, 2002). Particularly controversial rules are the abundant-centre and abundant niche-centre hypotheses, which predict abundance to decrease gradually from the centre to the margins of species geographic ranges and ecological niches respectively (Brown, 1984; Pironon *et al*., 2017). Both theories have received mixed empirical support (Martínez-Meyer *et al*., 2013; Sagarin & Gaines, 2002; Dallas *et al*., 2017) and limited theoretical development (Osorio-Olvera *et al*., 2019; Holt, 2019; Dallas & Santini, 2020). Moreover, recent analyses highlighted that tests of these hypotheses were sensitive to the quality of the input data and the methodological approach considered (Santini *et al*., 2019).

Osorio-Olvera *et al*. (2020) analyze data from the North American Breeding Bird Survey (BBS) to test for a negative correlation between species abundance and the distance to their climatic niche centroid. Counter to recent findings questioning its generalizability (Sagarin & Gaines, 2002; Dallas *et al*., 2017; Santini *et al*., 2019), the authors claimed general support for the hypothesis and proposed that the distance to species climatic niche centroid (quantified using minimum volume ellipsoids) could represent a reliable and simple new metric to predict the current and future distribution of species abundance. However, we discuss how serious problems related to data quality, modelling choice, and presentation of the results, prevent from making any reliable conclusion, and can greatly affect the perceived support for the hypothesis.

First of all, many of the species considered in the study also occur well beyond the study area (e.g. *Ardea alba, Corvus corax*), and some only share a very small portion of the range in the study area (e.g. *Thalasseus maximus, Aramus guarauna*). We calculated geographic and climatic niche overlap of a convex hull encompassing the BBS data with the BirdLife International data (BirdLife International, 2017) only considering the resident and breeding range, demonstrating a clear influence on the estimation of the geographic range and climatic niche boundaries, as well as their centroid distance (Figure 1). This subsequently affects the abundant niche-centre relationship, as discussed in Soberón *et al*. (2018), questioning the validity of the relationships estimated. Oddly, many of the bird species whose geographic ranges are underestimated and whose niches have been underestimated also exhibit significant negative abundant-centre relationships (e.g. *Tyrannus couchii, Thalasseus maximus, Glaucidium gnoma*), putatively supporting the hypothesis.

**Figure 1:**
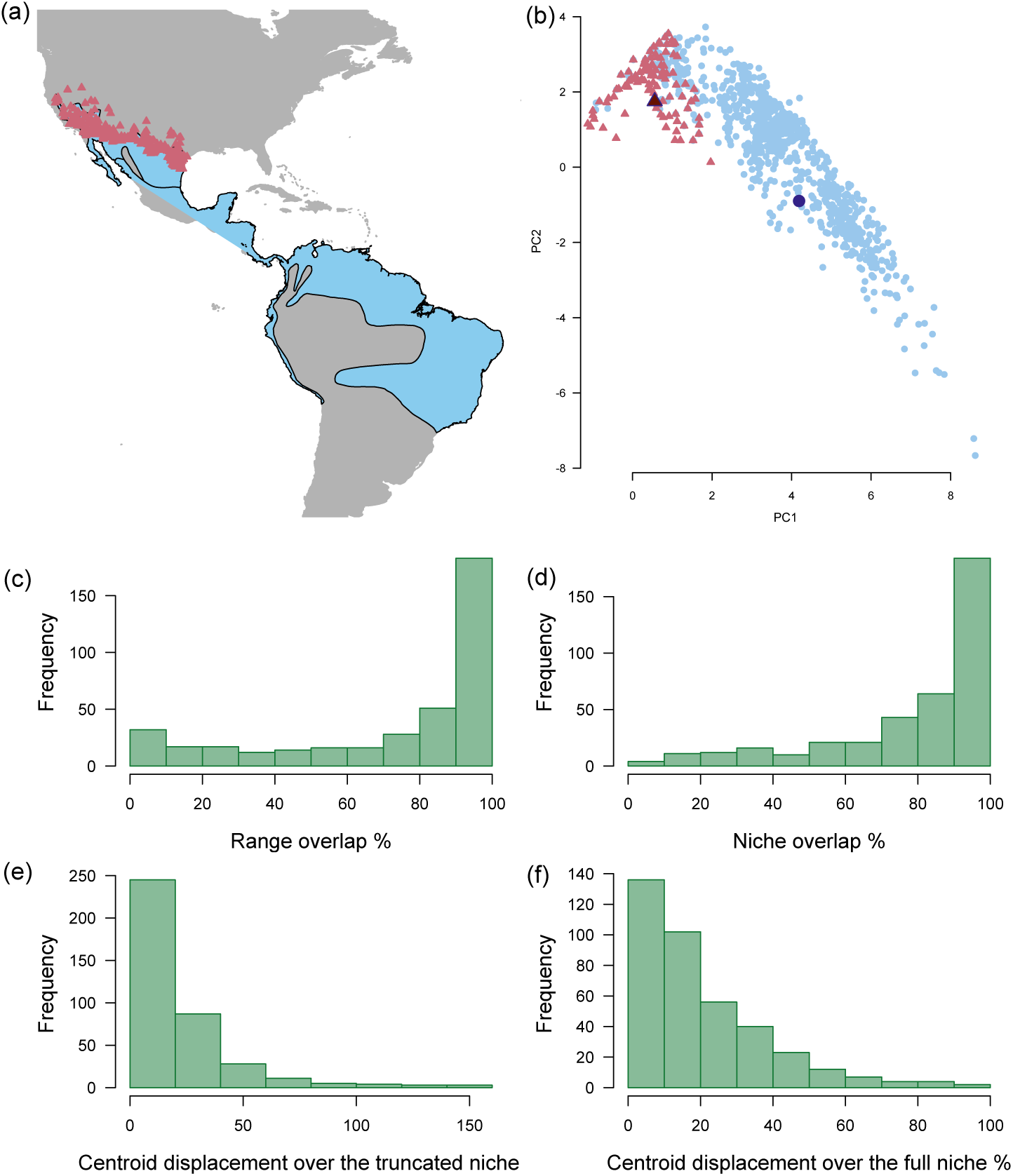
Mismatch in geographic and niche estimates between abundance data used in Osorio-Olvera *et al*. (2020) and the combined resident and breeding range species distributions, estimated IUCN range polygons. **a**) Lesser nighthawk (*Chordeiles acutipennis*) IUCN geographic range (in blue) and sample data to estimate the niche (in red); **b**) First two PCA axes of all bioclimatic variables showing environmental values considered in the study (red triangles) and those estimated considering the cells in the IUCN range (blue dots). The darker and larger triangle and circle represent the estimated centroids of the two hypervolumes; **c**) Distribution of geographic range overlap between convex hulls drawn around abundance estimates and the IUCN ranges for all species considered in the study; **d**) Distribution of niche overlap between convex hulls drawn around abundance estimates and grid cells within the IUCN ranges for all species in the study; **e**) Percentage of centroid displacement over the truncated niche; **f**) Percentage of centroid displacement over the full niche. Niche overlap and niche centroids were estimated using the hypervolume package Blonder *et al*. (2015).

The authors found that the percentage of species range overlap (Table 1 in Osorio-Olvera *et al*. (2020)) had a non-significant effect on the correlation coefficients (multivariate analysis), thus arguing that range overlap does not affect their conclusions. However, this may not account for the effect of niche truncation, as 1) geographic overlap does not necessarily translate into niche overlap (Fig. 1c,d), which is also why the authors estimate niche centres instead of geographic centres, and 2) the location of niche centres are still biased towards climatic conditions of the study area, which affects the calculation of centroids and distances (Fig. 1b). On a more fundamental level, testing if abundant-centre relationships differ as a function of range overlap does not address the influence of range overlap directly, but makes the assumption that as long as correlation coefficients do not differ as a function of range overlap, then the range centroid distances were estimated appropriately. This is not a clear test of the influence of range overlap, and risks the fallacy of *asserting the null*. We note that a biased estimation of the niche centre is supposed to matter in such an analysis, a non-significant difference suggests that using high-quality data does not increase the support rate for the hypothesis.

The strongest support for the abundant niche-centre relationships comes from Osorio-Olvera *et al*. (2020) estimating the species niche as a minimum volume ellipsoid (MVE) by considering more than 4000 combinations of climatic variables, including all 19 commonly-used bioclimatic variables together with the first 15 PCA components of a PCA based on the same bioclimatic variables. The authors use every possible combination of two and three niche axes to estimate the niche. We identify two main issues associated with this procedure.

First, the authors report results only for models showing significant abundant niche-centre relationships, omitting non-significant correlations (Figure 2a). This issue is not only present in the fit MVE models, but also in the 2 and 3 feature models using convex hulls or MVEs. The effects of this are clear (Figure 2). By including non-significant correlations, the mean abundant niche-centre relationship across all model sets becomes weak 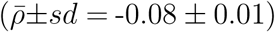, and more species exhibit significantly positive abundant niche-centre relationships (Figure 2). Including these non-significant results is important, in our view, and strongly influences the resulting perceived support for the abundant niche-centre pattern (Figure 2). Presenting also non-significant results demonstrates that only between 37% and 45% of species have negative abundant-centre relationships, regardless of approach used (see https://figshare.com/s/8fadf780810e73d44623), while the majority of the estimated relationships are either positive or non-significant. Interestingly, this low empirical support is consistent with previous findings (Dallas *et al*., 2017; Pironon *et al*., 2017; Sagarin & Gaines, 2002; Santini *et al*., 2019).

**Figure 2:**
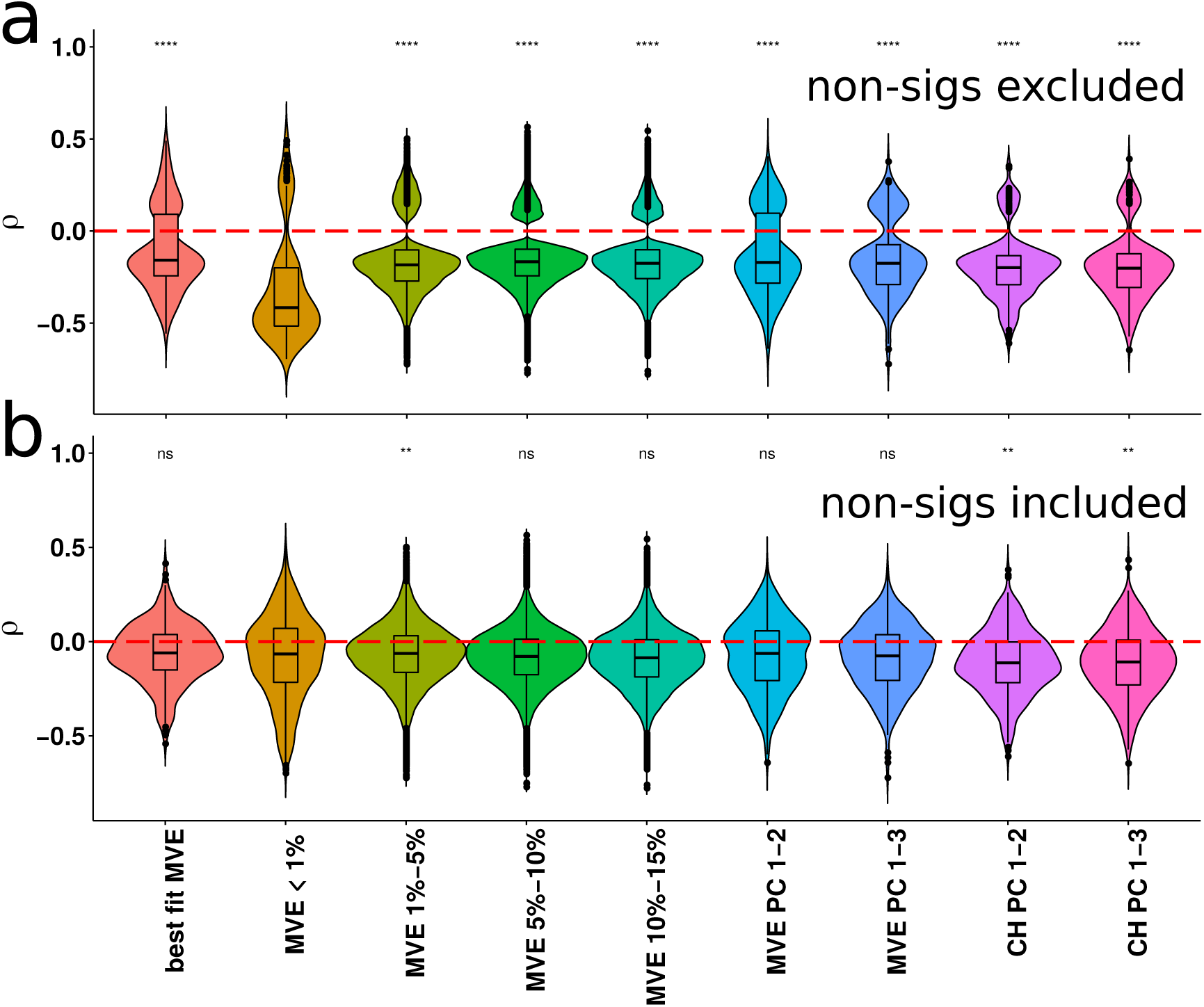
Differences in support for the abundant niche-centre hypothesis for a set of North American birds as a function of analytical decisions. We reproduce the results of Osorio-Olvera *et al*. (2020), demonstrating largely significant and negative abundant-centre relationships (panel a). However, by including all correlation coefficients, instead of only the significant ones, support for abundant niche-centre relationships become weak and largely non-significant (panel b). We also include the results when only considering the best fit MVE models per species (’best fit MVE’) when non-significant relationships were excluded (panel a) and included (panel b). Significance values (**** p *≤* 0.0001, *** p *≤* 0.001, ** p *≤* 0.01, * p *≤* 0.05, ns not significant), compare all other methods to the MVE *<* 1% case.

Second, while the authors train an average of 1,852 models per species to calculate MVEs, they perform no form of model selection (i.e., excluding models based on omission rate is thresholding, not model selection). This functionally treats the poorest fit MVE and the best fit MVE per species as equivalent, provided the model produced a significant abundant-centre relationship. This condition results in between 1 and 3460 abundant-centre estimates for any given species, introducing substantial bias in estimation of the distribution of abundant-centre relationships. When non-significant results are included, and only best models are retained, the overall pattern changes substantially (Figure 2). When only the best fit models are considered, 115 out of 379 species (30%) had significant abundant niche-centre relationships, with a mean correlation coefficient of -0.07. Some of these best models had higher omission rates than what Osorio-Olvera *et al*. (2020) considered. Removing these models reduces the number of species down to 303 species, of which 94 had significantly negative abundant niche-centre relationships (31%), while 180 and 29 had non-significant (59%) or significantly positive (10%) relationships, respectively (Figure 2c). It is not our assertion that abundant-centre relationships do not exist. The negative relationships found by Osorio-Olvera *et al*. (2020) support the idea of an abundant-centre, but do so in a misleading manner.

The study from Osorio-Olvera *et al*. (2020) highlights the timely need for disentangling the complex relationship between species ecological niche, geographic distribution and demographic performance (Holt, 2019; Bohner & Diez, 2020). Explaining the convergence and divergence of results of studies exploring occurrence and abundance patterns is key for improving our understanding of biodiversity and ability to predict its response to ongoing changes in the global environment.

## Data accessibility

R code is available on figshare at https://figshare.com/s/8fadf780810e73d44623.

